# Corticopostural functional and effective connectivity reveal cortical control of postural sway velocity during quiet standing

**DOI:** 10.1101/2023.12.14.571767

**Authors:** Maxime Fauvet, Clara Ziane, Ludovic Archambault-Levesque, Théo Fornerone, Fabien Dal Maso

## Abstract

**Background:** Despite a large body of evidence showing the involvement of the sensorimotor cortex in postural control, its exact role remains unclear. Models of postural control outcomes suggested that the velocity of the center of pressure is a crucial parameter to maintain balance. Inspired by corticokinematic coherence, we hypothesized that cortical oscillations and the velocity of the center of pressure (CoP) would synchronize and that this synchronization would increase with postural task difficulty during quiet standing.

**Methods:** We compared the magnitude of coherence and Granger causality computed between brain oscillations recorded with electroencephalography and the center of pressure velocity in the Delta and Theta frequency bands obtained from 23 participants performing four quiet standing tasks with various levels of difficulty. The effect of postural task difficulty and information flow direction were tested with a linear mixed model while non-parametric correlations were computed between coherence magnitude and postural performance measured by 95% confidence ellipse area and mean center of pressure velocity.

**Results:** We found significant coherence between the Cz EEG electrode and CoP velocity in the Delta and Theta frequency bands. This EEG-CoP velocity coherence significantly increased with task difficulty in the Delta (F = 18.8, p < 0.001) and Theta (F = 7.83, p < 0.001) bands. Granger causality significantly increased with task difficulty (F = 12.5, p < 0.001) and was higher in the efferent than afferent direction (F = 78, p < 0.001). The 95% confidence ellipse area was correlated to coherence magnitude in the most difficult condition. Participants showing significant Granger causality in the afferent direction showed more stable postural outcomes.

**Conclusion:** Our results confirm that the CoP velocity has a crucial role in postural control through its synchronization with sensorimotor cortex oscillations. The efferent information predominance suggests that posture is partly controlled by the sensorimotor cortex by a mechanism named corticopostural coherence. Our results show that this corticopostural coherence could represent a mechanism for controlling balance during quiet standing.

## 1 Introduction

Understanding postural control mechanisms, particularly during quiet standing, is crucial for improving the quality of life of the growing elderly population since falls pose a significant threat to their well-being and independence (Kalache et al., 2007). For a long time, quiet standing was believed to be controlled by subcortical and peripheral mechanisms (Magnus, 1926; Sherrington, 1910), due to its automatic and reflexive apparent nature (Varghese et al., 2014). However, a pioneering research employing a dual-task paradigm combining a balance task with a memory load (Kerr et al., 1985), revealed a competition between visuospatial memory and postural performance that was interpreted as an indirect evidence of cortical involvement in postural control. Subsequent studies supported this interpretation reporting postural adjustments to imbalance occurred at varying latencies, with shorter latencies involving spinal or subcortical loops (Ackerman, 1991), and longer latencies involving cortical networks (e.g., Taube et al., 2006). Despite a considerable body of research evidencing cortical involvement in postural control (Hülsdünker et al., 2015; Jacobs & Horak, 2007; Mihara et al., 2008; Slobounov et al., 2005; Stokkermans et al., 2023; Varghese et al., 2014), the precise role of the sensorimotor cortex in quiet standing remains poorly understood.

Much of our current understanding regarding the cortical role in postural control stems from electroencephalography (EEG) studies conducted during postural tasks. For instance, Slobounov et al. (2005) found Delta (1-4 Hz) transient activity in the central sensorimotor cortex during self-initiated swings. Additionally, other studies have consistently identified a negative event-related potential in the fronto-central area 100 ms after a balance perturbation (Jacobs & Horak, 2007; Marlin et al., 2014). Using source reconstruction, Theta frequency band (4-8Hz) power was shown to increase in the anterior and posterior cingulate cortex in response to balance loss (Sipp et al., 2013). These patterns are suggested to represent the integration of sensory information in different brain regions required to maintain balance after a perturbation (Burgess et al., 1997). Aside from adaptation to balance perturbations, investigations also focused on quiet standing, a simple paradigm for studying postural control at both central and peripheral levels (Winter, 1995). Studies have shown an augmented involvement of cortical activity during postural tasks of increasing difficulty. For example, Hülsdünker et al. (2015) found an increased Theta power spectrum in central and parietal regions and an increase in Alpha peak frequency during a postural task of increasing difficulty (Hülsdünker et al., 2016). Alpha and Beta desynchronization as well as Delta and Theta synchronization were found to increase in the central area concurrently with task difficulty (Edwards et al., 2018; Liang et al., 2022; Ozdemir et al., 2018). The latter results were interpreted as a heightened afferent information processing. However, despite the wealth of information about how cortical activity is affected by various balance conditions, we still lack a comprehensive explanation of the cortical oscillations’ role in postural control.

Postural control during quiet standing is commonly described through the Single Inverted Pendulum paradigm, where the center of mass (CoM) of the body is maintained within the base of support while the body sways in the anteroposterior and mediolateral directions (Morasso et al., 2019). Originally, based on Feldman’s theory of equilibrium point (Feldman, 1986), Winter (1995) proposed that the Single Inverted Pendulum is controlled at the ankle joint through passive muscle stiffness, requiring minimal involvement from the central nervous system. However, subsequent studies showed that ankle muscle stiffness alone was not sufficient to maintain equilibrium (Casadio et al., 2005; Kiemel et al., 2002; Loram & Lakie, 2002; Soest et al., 2003; van der Kooij et al., 2001). As a result, the stiffness model was expanded to include an active control requiring the involvement of the central nervous system (Kiemel et al., 2002; Mergner et al., 2002; van der Kooij et al., 2001), with Morasso et al. (2019) proposing a framework based on intermittent active control (Asai et al., 2009; Bottaro et al., 2008). In the latter model, the velocity of the CoM emerged as a variable controlled by the central nervous system. The body sway velocity was considered more efficient to control posture than the position of the CoM, as it provides valuable inferences about the system’s future state (Asai et al., 2009; Delignières et al., 2011). Masani et al. (2003) showed a cross-correlation between ankle muscles electromyography and CoM velocity, revealing that the pattern of ankle muscle activity precedes the pattern of CoM velocity. Furthermore, selective alteration of the CoM velocity feedback provided by vision and proprioception led to poorer postural performance (Jeka et al., 2004). CoM and center of pressure (CoP) and their related outcomes have been largely used in characterizing quiet standing (Richmond et al., 2021). Interestingly, CoM movement can be seen as the origin of imbalance, while the CoP reflects the response of the central nervous system to maintain the CoM within the base of support (Richmond et al., 2021). Collectively, these findings provide compelling evidence that body sway velocity is fundamental to control balance during quiet standing, but to date, no previous investigation explored the cortical mechanism underlying its control.

Previous studies on brain dynamics and body kinematics have evidenced significant coupling between cortical oscillations and movement velocity through coherence analyses (Bourguignon et al., 2019), the so-called corticokinematic coherence (CKC). Coherence is a measure of correlation in the frequency domain between oscillatory time series (Rosenberg et al., 1989) with values ranging from 0 to 1 respectively reflecting no synchronization and perfect synchronization between oscillatory components. Coherence has been extensively investigated between electrophysiological signals as a means of assessing large-scale integration and functional connectivity between remote population of neurons (Bastos & Schoffelen, 2016; Varela et al., 2001). By employing CKC analysis, Jerbi et al. (2007) were the first to evidence significant synchronization between cortical oscillations from the primary motor cortex and hand velocity at low frequencies (2-5 Hz) during a track-ball manipulation task. Additional studies revealed synchronization at the movement frequency between cortical oscillations from the sensorimotor area and finger acceleration during rhythmic movements (Bourguignon et al., 2011, 2012). When investigating the direction of information flow via Granger causality analysis (GC), authors found that CKC was mainly driven by afferent information to the brain, reflecting sensorimotor integration of movement velocity by the primary somatosensory area (Bourguignon et al., 2015; Piitulainen et al., 2013). To the best of our knowledge, no studies have explored the synchronization between cortical oscillations and body sway velocity during postural tasks, although the later seems to have a critical role in quiet standing control as previously introduced.

Here, we aimed to enhance our understanding of the sensorimotor cortex’s involvement in postural control. To this end, we performed spectral power, coherence and information flow direction analyses between cortical and CoP velocity oscillations during postural tasks with different levels of difficulty. CoP was selected over CoM because it thought to better reflect the responses of the central nervous system to unbalance (Richmond et al., 2021). We hypothesized that postural tasks of increasing difficulty would induce a desynchronization in Alpha and Beta bands in central and parietal areas (Hülsdünker et al., 2015, 2016; Liang et al., 2022) as well as a central synchronization in Theta and Delta bands (Edwards et al., 2018; Ozdemir et al., 2018). Drawing on previous CKC findings (Jerbi et al., 2007), the importance of CoM velocity in postural control (Masani et al., 2003), and in accordance with the sensorimotor cortex involvement during postural control (Jacobs & Horak, 2007), we expected evidencing significant coherence between cortical and CoP velocity oscillations in low frequency band. We also expected that this EEG-CoP velocity coherence would increase over central cortical area with task difficulty. In addition, we expected a positive correlation between postural outcomes and cortical spectral power and coherence. Finally, we anticipated finding a higher afferent than efferent GC (Bourguignon et al., 2019), and that this difference would increase with postural task difficulty (Hülsdünker et al., 2016).

## 2 Methods

### 2.1 Participants

Twenty-eight young adult right footed participants (14 females, 25.4 ± 3.9 years old, height = 172. ± 7.7 cm, weight = 65.5 ± 8.3 kg) were recruited. Participants with any of the following conditions were not included, namely, body mass index over 30, neurological disorders, or recent physical injuries that could impair balance control. The Research Ethics Committee of the Université de Montréal (Montréal, Canada, CERC2019-27) approved the experimental protocol. All participants provided their written informed consent before any experimental procedure took place.

### 2.2 Data acquisition

Ground reaction forces and moments were recorded at 100 Hz using a six-axis (Fx, Fy, Fz and Mx, My, Mz) BP900900 force platform (AMTI, Watertown, Massachusetts, USA). EEG signals were recorded at 1000 Hz using 64 ActiChamp slim electrodes (Brain Product, Munich, Germany) located on the EEG cap according to the 10-20 system. The ground electrode was located at the AFz electrode location, and all electrode signals were online referenced to the FCz electrode. Electrode impedances were kept below 20 kOmhs with a conductive gel (SuperVisc, Easycap). Force platform and EEG signals were synchronized offline using a Transistor-Transistor Logic signal.

### 2.3 Experimental protocol

Participants performed four quiet standing experimental conditions that combined feet stance width and ground compliance to modulate postural balance. Ranging from the easiest to the most challenging postural task (Liang et al., 2022; Patel et al., 2008), participants stood upright in four conditions combining Normal (feet at shoulder width) and Narrow (feet as close as possible without touching) stances with different surface compliances, including Ground (force platform) and Foam (10-cm medium-density foam). In all experimental conditions, participants wore their socks to prevent surface temperature changes between the Ground and Foam conditions. They were instructed to stand upright with their arms resting alongside their body and keep their eyes open. Additionally, a resting state trial was recorded while participants sat on a chair to compute task-related spectral perturbation (TRSP, see below for details). All experimental conditions were recorded for a total of 300 seconds, separated in two 150 seconds trials to avoid fatigue. All trials were randomized across participants.

### 2.4 EEG signal pre-processing

EEG data pre-processing was performed offline using the EEGLAB Matlab toolbox (Delorme & Makeig, 2004) and custom-made Matlab scripts (Matlab 2017b version). The raw EEG signals were bandpass filtered between 1 and 200 Hz and stopband filtered at 60, 120, and 180 Hz (±0.25 Hz around each frequency) to remove line noise. Both filters were zero-lag second order Butterworth filters. Thereafter, EEG channels with an excessive level of noise were automatically rejected using the 2.7 version of the *Clean RawData plugin* with ‘flat line criterion’ set to 5 seconds and ‘minimum channel correlation’ set to 0.6. Moreover, EEG channels presenting standard deviation superior to 1000 µV or a kurtosis superior to five times the mean were removed (Gwin et al., 2011). On average (std), 61.1 (5.8) EEG channels were kept for subsequent analyses. The two trials of each experimental condition were concatenated. EEG signals were then re-referenced to a common average reference. The Infomax algorithm was used to decompose independent components (ICs) from the EEG signals and resulting ICs were labelled using the ICLabel plugin (Pion-Tonachini et al., 2019). Labelled ICs containing a probability of muscle, eye, heart, line noise, channel noise, and other above 50%, 50%, 35%, 35%, 35%, and 80%, respectively, were rejected. On average (std), 36.3 (5.1) ICs were rejected. Finally, EEG signals and ICs were visually inspected for any remaining source of contamination. The data from three participants were removed from further analysis due to poor quality of EEG signals. The data of two additional participants were removed due to inconsistent force platform signals, leading to a final sample of 23 participants.

### 2.5 Data analysis

#### 2.5.1 Postural data

Force platform signals were lowpass filtered at 20 Hz with a zero-lag fourth order Butterworth filter. CoP position time series in the anteroposterior and mediolateral directions were used to compute the 95% confidence ellipse area and the mean CoP velocity. The 95% confidence ellipse area is a statistical representation of the average CoP displacement. Its increase has been associated with an increased risk of falls (Merlo et al., 2012). The mean velocity of the CoP is a dynamic variable computed from the total sway length divided by the trial duration. An increase in mean velocity has been associated with risks of falls (Howcroft et al., 2017) and postural imbalance (Prieto et al., 1996; Prieto & Myklebust, 1993). Finally, power spectrum of the CoP velocity was obtained using a Short Time Fourier Transform algorithm. Frequency power increases in the 0-2 Hz frequency band have been associated with diminished stability in healthy young adults (Sozzi et al., 2021).

#### 2.5.2 Spectral power analysis of EEG signals

Pre-processed EEG signals were time-frequency transformed using the *wavelet* Matlab function (Torrence & Compo, 1998) with the following settings: Morlet wavelet as mother wavelet and 7 as wavenumber, yielding 161 evenly spaced scales between 0 and 50 Hz. The spectral power (POW) of Delta (1-4 Hz), Theta (4-8 Hz), Alpha (8-13 Hz), and Beta (13-30 Hz) frequency bands were computed from the wavelet power spectrum, averaged over the total duration of each condition and normalized to the resting state trial using the following equation to obtain the TRSP (Pfurtscheller & Aranibar, 1977):

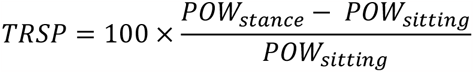

Negative and positive TRSP values indicate desynchronization and synchronization, respectively.

#### 2.5.3 Coherence between cortical and CoP velocity oscillations

##### Coherence analysis

The coupling between the oscillations of the CoP velocity and all EEG electrodes was estimated using coherence analysis. Prior to the analysis, EEG signals and COP velocity were epoched into 10-second epochs. Ten-second epoch durations were chosen to have a sufficient number of data segments (*i*.*e*., 30) to achieve a correct coherence estimation, while having an accurate time-frequency estimation of the lowest frequency components (*i*.*e*., Delta frequency band). As the EEG-CoP velocity coherence is expected in the Delta and Theta frequency bands and the power spectrum components of CoP velocity does not exceed 10 Hz (Demura et al., 2008), we adapted the parameters of the *wavelet* function to improve the low frequency components identification. Thus, epoched signals were time-frequency transformed using the *wavelet* Matlab function using a wavenumber of 4 and thus yielding 92 evenly spaced scales between 0 and 20 Hz. Magnitude squared coherence (MSCoh) was computed for all frequencies between 0 and 20 Hz according to the following equation:

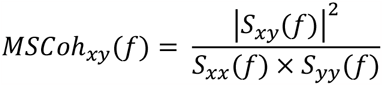

where *S*_*xy*_ is the crosspectrum between CoP velocity and the EEG signal and *S*_*xx*_ and *S*_*yy*_ are the autospectra of CoP velocity and the EEG signal, respectively (Bastos & Schoffelen, 2016). The first level significance of the EEG-CoP velocity coherence was computed for each experimental condition on time-frequency representations using the 95% confidence threshold *L* = 1 − (α)^1/(*n*−1)^, where α is 0.05 and n is the number of epochs used for the coherence calculation (Rosenberg et al., 1989). Coherence magnitude in the Delta and Theta frequency bands was obtained by averaging the EEG-CoP velocity coherence values above the significant threshold over time and frequency.

##### Granger causality analysis

The previous coherence analysis was completed with a GC analysis in order to identify the direction of the information flow between the EEG and the CoP velocity oscillations. Given the high computational cost of GC analyses, signals were downsampled to 100 Hz. Additionally, only the Cz electrode was analysed based on the EEG-CoP velocity coherence results. The GC analysis was performed with the Multi Variate Granger Causality toolbox (Barnett & Seth, 2014). A model order of 9 was chosen according to the Akaike Information Criterion and Bayes Information Criterion averaged over all participants, yielding an average model prediction window of 90 ms in both directions, *i*.*e*., EEG preceding CoP (EEG->CoP) and CoP preceding EEG (CoP->EEG).

### 2.6 Statistical analysis

All statistical analyses were performed with the version 2.3 Jamovi (The jamovi project (2023)) and R (v4.2.1; R Core Team, 2022). The effect of task difficulty on the area of the 95% confidence ellipse, the mean CoP velocity, the TRSP and EEG-CoP velocity coherence were analysed through linear mixed models with ‘task difficulty’ as a four-level fixed factor (Normal_Ground_, Narrow_Ground_, Normal_Foam_, Narrow_Foam_), and ‘participants’ as random factor: *Variable*∼ 1 + *task*_*difficulty* + (1|*participants*). The GC was analysed with a linear mixed model with two fixed factors ‘task difficulty’ (four levels: Normal_Ground_, Narrow_Ground_, Normal_Foam_, Narrow_Foam_) and ‘flow_direction’ (two levels: EEG->CoP, CoP->EEG), the interaction between these factors, and ‘participants’ as random factor: *GC* = 1 + *task*_*difficulty* ∷ *flow_durection* + (1|*participants*). Pairwise t-tests with Bonferroni corrections were applied as post-hoc tests when necessary and pairwise comparisons in TRSP and multiple comparisons of EEG-CoP velocity coherence were corrected using false discovery rate (FDR) (Benjamini & Hochberg, 1995). Models’ residuals were visually inspected with Q-Q plots to assess their normal distribution and Cook’s distance plots to evaluate the most influential points are provided in supplementary materials. All model results are presented with r-squared estimates, F or t values, degrees of freedom, 95% confidence intervals, and p-values.

Due to non-normality of TRSP, EEG-CoP velocity coherence, and postural outcomes distributions, Spearman’s rank correlations were performed to test the correlation between electrophysiological signals and postural outcomes. Correlation p-values were Holm adjusted and both ρ and p-values are reported.

## 3 Results

### 3.1 Postural outcomes

#### 95% confidence ellipse area

The linear model accounted for 27% of the total variance (Figure 1). The area of the 95% confidence ellipse increased with task difficulty (F_66_ = 17, p <.001) with an increase in area between Normal_Ground_ and Narrow_Foam_ conditions comprised in the 95% confidence interval [1392-2516] cm^2^. Post-hoc tests results are reported in Table 1. *Mean CoP velocity*. The linear model accounted for 70% of the total variance (Figure 1). The mean CoP velocity increased with task difficulty (F_66_ = 231, p < .001) with an increase between Normal_Ground_ and Narrow_Foam_ conditions comprised in the 95% confidence interval [0.01146 - 0.01345] cm/s. The post-hoc tests showed significant differences (p < .001) in the mean CoP velocity between all experimental conditions (Table 1). The residuals of both 95% confidence ellipse area and mean CoP velocity linear mixed models were normally distributed, yielding a very high confidence in models’ results.

**Figure 1:**
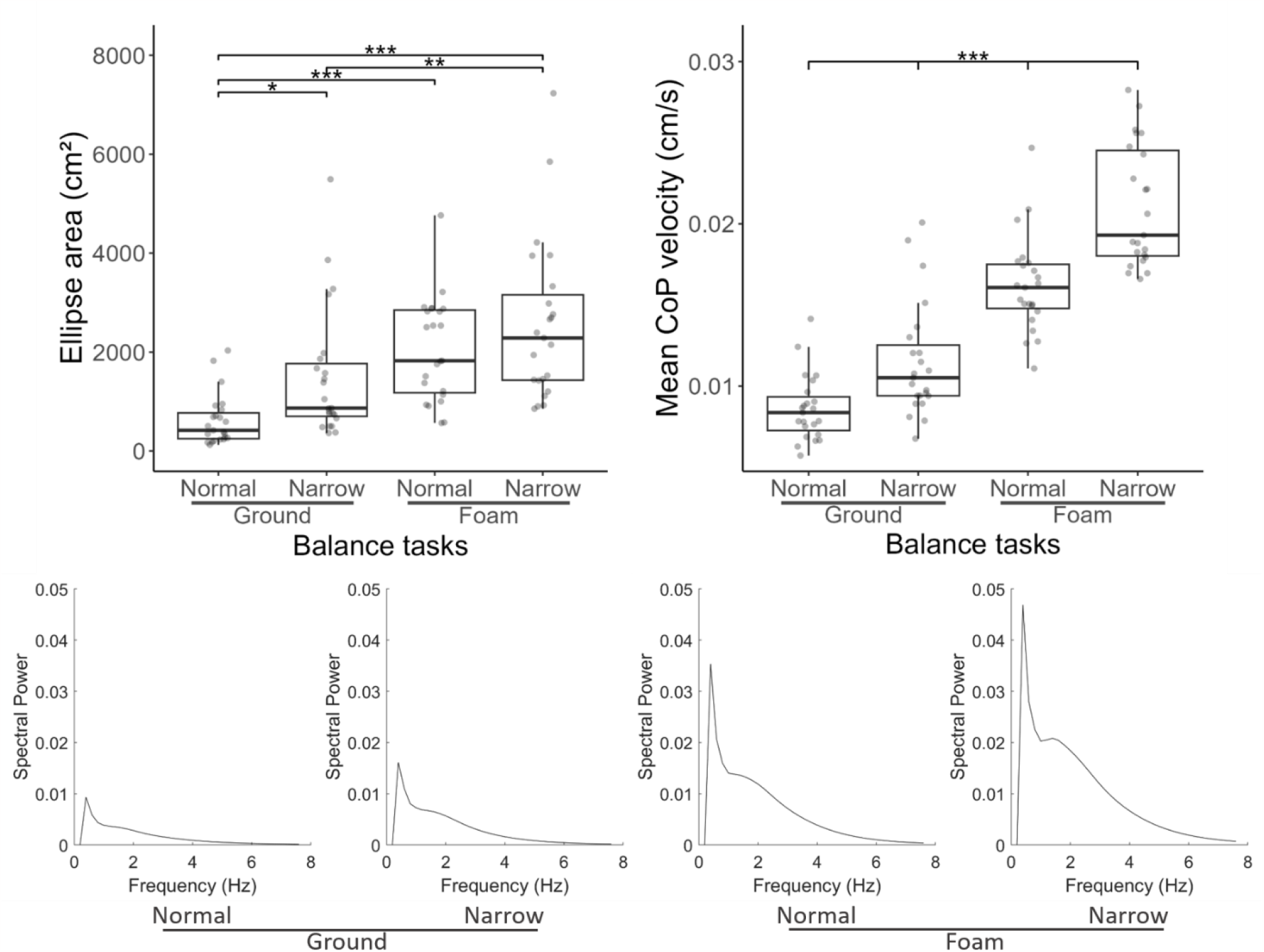
Postural outcomes in each experimental condition. Top row: 95% confidence ellipse area (left) and mean CoP velocity (right). Dots in boxplots represent participants’ individual values. Bottom row: Group average power spectra of the CoP velocity in each experimental condition. *p < .05, **p < .01, ***p < .001.

**Table 1:** Post-hoc results of 95% confidence ellipse area and mean CoP velocity analyses.

| Comparison |  | 95% confidence ellipse area |  |  | Mean CoP velocity |  |  |
| --- | --- | --- | --- | --- | --- | --- | --- |
| Task difficulty | Task difficulty | Estimate (cm²) | t | p<sub>bonferroni</sub> | Estimate (cm/s) | t | p<sub>bonferroni</sub> |
| Normal<sub>Ground</sub> | Narrow<sub>Ground</sub> | -875 | -3.05 | 0.020 | -0.00288 | -5.66 | <.001 |
| Normal<sub>Ground</sub> | Normal<sub>Foam</sub> | -1438 | -5.01 | <.001 | -0.00760 | -14.94 | <.001 |
| Normal<sub>Ground</sub> | Narrow<sub>Foam</sub> | -1954 | -6.81 | <.001 | -0.01246 | -24.48 | <.001 |
| Narrow<sub>Ground</sub> | Normal<sub>Foam</sub> | -562 | -1.96 | 0.326 | -0.00472 | -9.28 | <.001 |
| Narrow<sub>Ground</sub> | Narrow<sub>Foam</sub> | -1079 | -3.76 | 0.002 | -0.00958 | -18.82 | <.001 |
| Normal<sub>Foam</sub> | Narrow<sub>Foam</sub> | -517 | -1.80 | 0.458 | -0.00485 | -9.54 | <.001 |

The group-averaged power spectra of the CoP velocity showed a peak at 0.5 Hz (Figure 1). 99% of the total power spectra was comprised in the 0-8 Hz frequency band in each experimental condition. 52%, 55%, 55%, and 57% of the total power spectra was comprised in the Delta frequency band and 13%, 11%, 12%, and 15% was comprised in the Theta frequency band in the Normal_Ground_, Narrow_Ground_, Normal_Foam_ and Narrow_Foam_ conditions, respectively.

### 3.2 Task-Related Spectral Perturbation

Beta and Alpha frequency bands showed a desynchronization in the central and parietal areas (Figure 2). Theta and Delta bands showed a synchronization at the Cz electrode site in the Normal_Foam_ and Narrow_Foam_ experimental conditions. No significant differences in spectral power were found between baseline and postural tasks (FDR corrected p-values > 0.05). In addition, no significant correlations were observed between the TRSP and the postural outcomes regardless of the frequency band or experimental condition.

**Figure 2:**
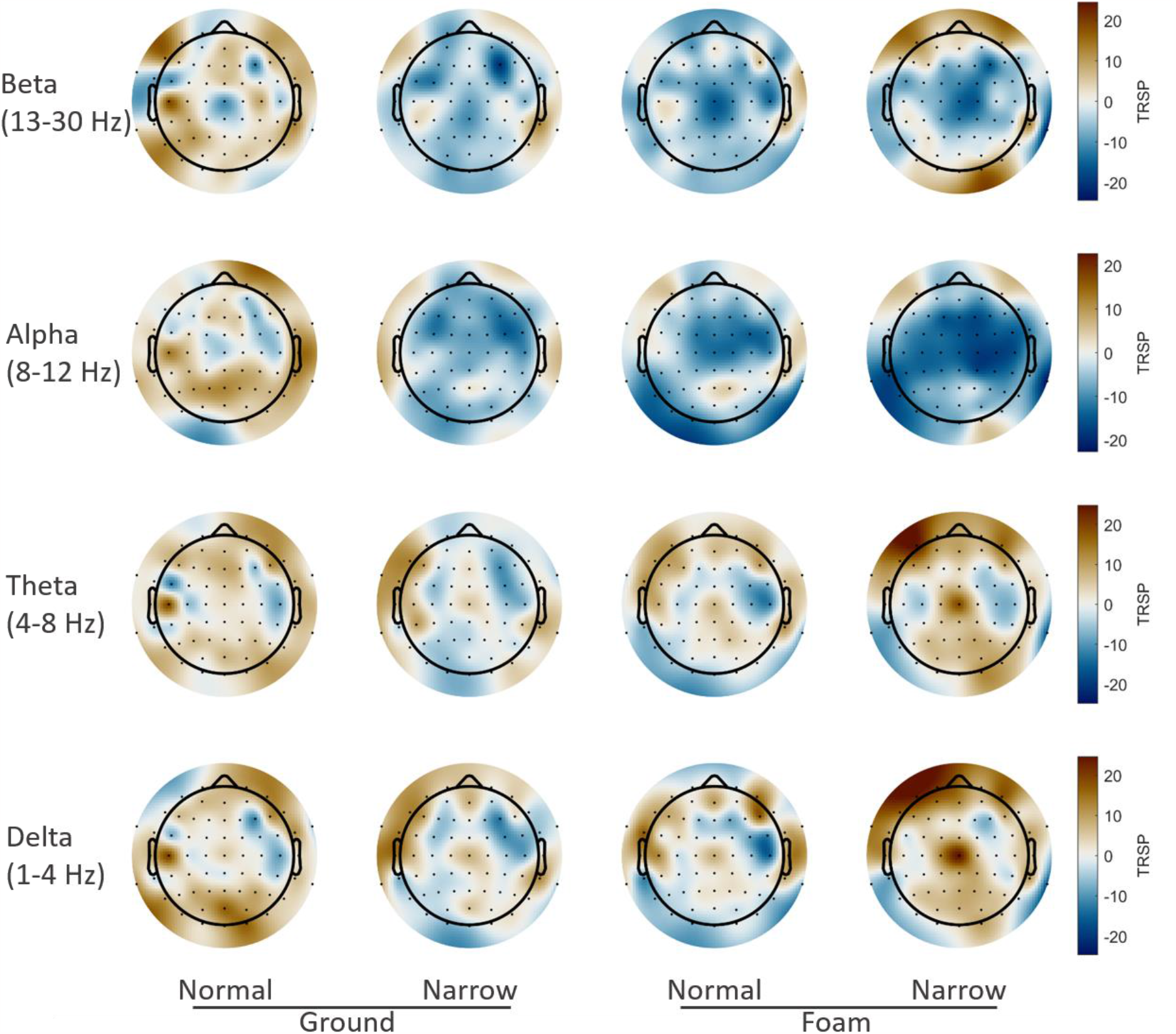
Group average TRSP topographies in each experimental condition in the Beta, Alpha, Theta, and Delta frequency bands (from top to bottom).

### 3.3 Coherence between cortical and CoP velocity oscillations

EEG-CoP velocity coherence increased at the Cz and neighbourhood electrodes with task difficulty, while other electrodes remained non-significant (FDR corrected p-values > 0.05) (Figure 3).

**Figure 3:**
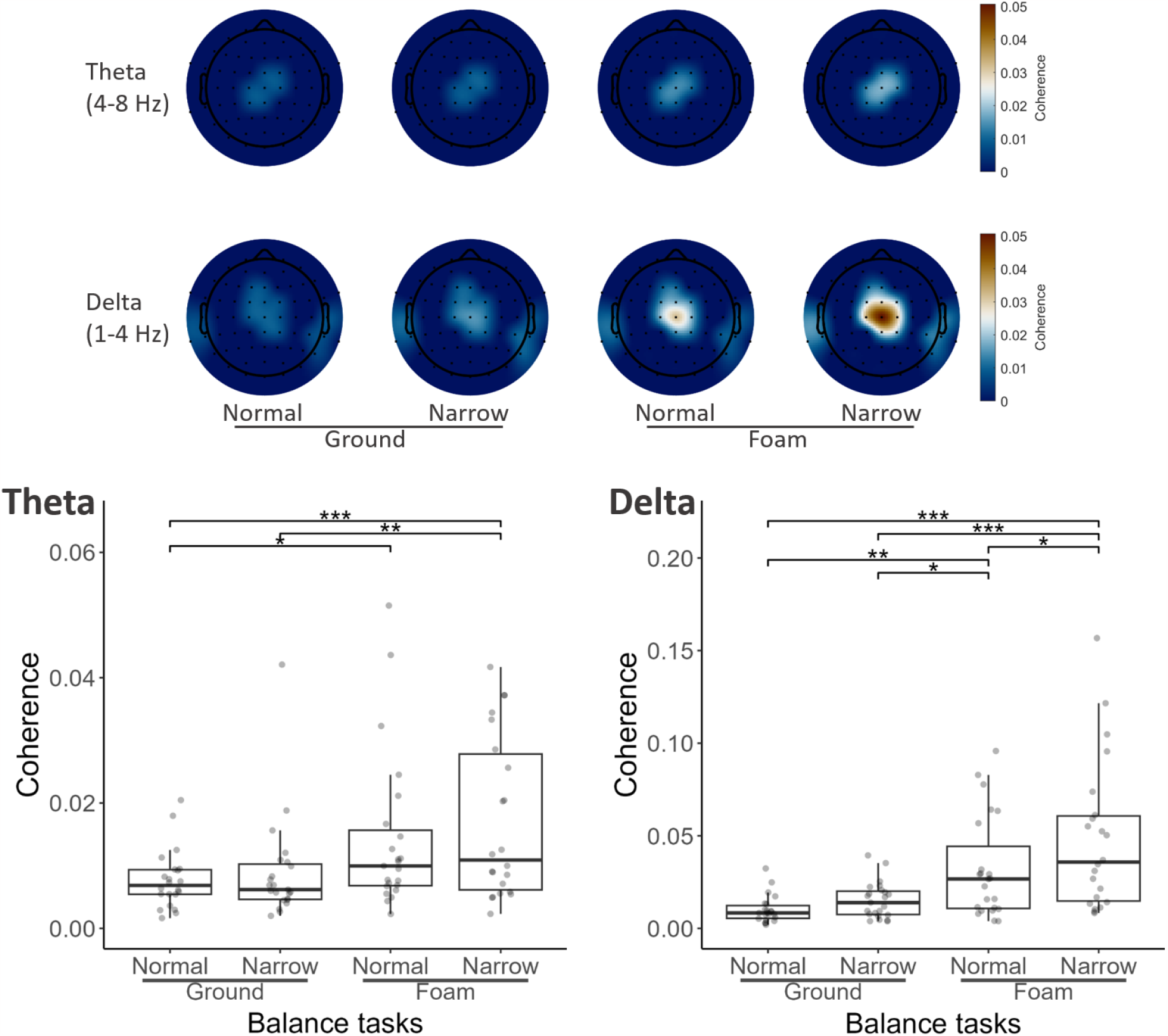
Group average topographies of Theta (top row) and Delta (middle row) frequency bands EEG-CoP velocity coherence in each experimental condition. Cz electrode EEG-CoP velocity coherence in Theta (lower row, left) and Delta (lower row, right) frequency bands in each experimental condition. Dots in boxplots represent participants’ individual values. Only first level significant EEG-CoP velocity coherence values were included in-group averages. *p < .05, **p < .01, ***p < .001.

#### Theta band

The linear model accounted for 13% of the total variance in Cz EEG-CoP velocity coherence (Figure 3). Theta band EEG-CoP velocity coherence increased with task difficulty (F_64_ = 7.83, p < .001) with an increase comprised in the 95% confidence interval [0.0051 – 0.0143] between Normal_Ground_ and Narrow_Foam_ conditions. *Delta band*. The linear model accounted for 27% of the total variance in Cz EEG-CoP velocity coherence (Figure 3). Delta band EEG-CoP velocity coherence increased with task difficulty (F_64_ = 18.1, p <.001) with an increase comprised in the 95% confidence interval [0.0058 – 0.027] between Normal_Ground_ and Narrow_Foam_ conditions. Post-hoc tests results are shown in Table 2 for both models. The residuals of both linear mixed models were mostly normally distributed, yielding a high confidence in models’ results.

**Table 2:** Post-hoc results of Theta and Delta EEG-CoP velocity coherence analyses at the Cz electrode.

| Comparison |  | Cz Theta coherence |  |  | Cz Delta coherence |  |  |
| --- | --- | --- | --- | --- | --- | --- | --- |
| Task difficulty | Task difficulty | Estimate | t | p<sub>bonferroni</sub> | Estimate | t | p<sub>bonferroni</sub> |
| Normal<sub>Ground</sub> | Narrow<sub>Ground</sub> | -0.00148 | -0.654 | 1 | -0.00557 | -0.967 | 1 |
| Normal<sub>Ground</sub> | Normal<sub>Foam</sub> | **-0.00687** | **-3.047** | **0.020** | **-0.02246** | **-3.897** | **0.001** |
| Normal<sub>Ground</sub> | Narrow<sub>Foam</sub> | **-0.00955** | **-4.176** | **<.001** | **-0.03861** | **-6.609** | **<.001** |
| Narrow<sub>Ground</sub> | Normal<sub>Foam</sub> | -0.00540 | -2.247 | 0.108 | **-0.01688** | **-2.972** | **0.025** |
| Narrow<sub>Ground</sub> | Narrow<sub>Foam</sub> | **-0.00808** | **-3.581** | **0.004** | **-0.03304** | **-5.734** | **<.001** |
| Normal<sub>Foam</sub> | Narrow<sub>Foam</sub> | -0.00268 | -1.188 | 1 | **-0.01616** | **-2.804** | **0.040** |

Cz Delta band EEG-CoP velocity coherence showed a significant negative correlation with the 95% confidence ellipse area in the Narrow_Foam_ condition (ρ = -0.45, p = 0.038). A significant negative correlation was found between the Cz EEG-CoP velocity coherence in the Theta band and the 95% confidence ellipse area in Narrow_Ground_ (ρ = -0.60, p = 0.003) and Narrow_Foam_ (ρ = -0.60, p = 0.004) conditions. These latter results indicate that the increase of the 95% confidence ellipse area was correlated to smaller Cz EEG-CoP velocity coherence. No significant correlations between the mean CoP velocity and the Cz EEG-CoP velocity coherence in the Delta or Theta bands were observed (Table 3).

**Table 3:**
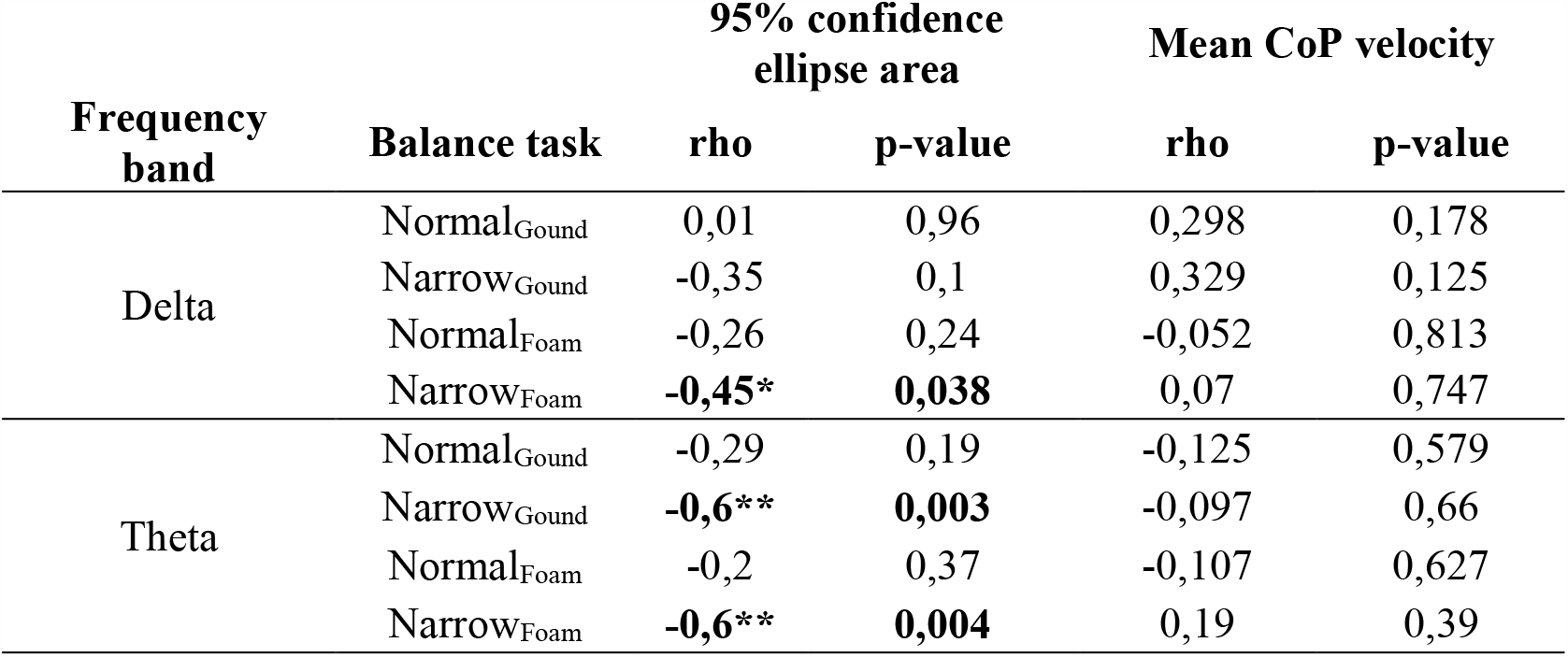
Correlation between the Cz EEG-CoP velocity coherence in the Delta and Theta frequency bands and 95% confidence ellipse area and mean CoP velocity. *p < .05, **p < .01.

### 3.4 Granger causality between cortical and CoP velocity oscillations

The linear mixed model accounted for 35% of the total variance in the GC and showed significant main effects of task difficulty (F_151_ = 12.47, p < .001) and information flow direction (F_150_ = 78.03, p < .001), and an interaction between task difficulty and information flow direction factors (F_150_ = 4.17, p = 0.007) (Figure 4, upper panel). The GC increased more with task difficulty in the EEG to CoP velocity direction flow than in the CoP velocity to EEG direction flow. The GC increased of 0.00212 (122% of increase, 95% confidence interval: [0.00130 – 0.00294]) between Normal_Ground_ and Narrow_Foam_ conditions and the slope of the direction of information flow effect was 0.00257 (95% confidence interval: [0.00200 – 0.00314]). The model’s residuals distribution was slightly different from a normal distribution, yielding a medium confidence in the models’ results and indicating skewness in the data. Post-hoc tests results are shown in Table 4.

**Table 4:** Post-hoc tests of the interaction term. Bold rows show significant differences between the direction flows for each task difficulty.

| Comparison |  |  |  |  |  |  |  |
| --- | --- | --- | --- | --- | --- | --- | --- |
| Task difficulty | Direction flow | vs. | Task difficulty | Direction flow | t | dof | Pbonferroni |
| Normal<sub>Ground</sub> | EEG → CoP | - | Normal<sub>Ground</sub> | CoP → EEG | 2.019 | 150 | 1 |
| Normal<sub>Ground</sub> | EEG → CoP | - | Narrow<sub>Ground</sub> | EEG → CoP | -1.657 | 150 | 1 |
| Normal<sub>Ground</sub> | EEG → CoP | - | Narrow<sub>Ground</sub> | CoP → EEG | 1.831 | 150 | 1 |
| Normal<sub>Ground</sub> | EEG → CoP | - | Normal<sub>Foam</sub> | EEG → CoP | -5.583 | 150 | < .001 |
| Normal<sub>Ground</sub> | EEG → CoP | - | Normal<sub>Foam</sub> | CoP → EEG | 1.025 | 150 | 1 |
| Normal<sub>Ground</sub> | EEG → CoP | - | Narrow<sub>Foam</sub> | EEG → CoP | -5.297 | 151 | < .001 |
| Normal<sub>Ground</sub> | EEG → CoP | - | Narrow<sub>Foam</sub> | CoP → EEG | 0.144 | 151 | 1 |
| Normal<sub>Ground</sub> | CoP → EEG | - | Narrow<sub>Ground</sub> | CoP → EEG | -0.208 | 150 | 1 |
| Normal<sub>Ground</sub> | CoP → EEG | - | Normal<sub>Foam</sub> | CoP → EEG | -1.014 | 150 | 1 |
| Normal<sub>Ground</sub> | CoP → EEG | - | Narrow<sub>Foam</sub> | CoP → EEG | -1.87 | 151 | 1 |
| Narrow<sub>Ground</sub> | EEG → CoP | - | Normal<sub>Ground</sub> | CoP → EEG | 3.696 | 150 | 0.009 |
| **Narrow<sub>Ground</sub>** | **EEG → CoP** | - | **Narrow<sub>Ground</sub>** | **CoP → EEG** | **3.533** | **150** | **0.015** |
| Narrow<sub>Ground</sub> | EEG → CoP | - | Normal<sub>Foam</sub> | EEG → CoP | -3.976 | 150 | 0.003 |
| Narrow<sub>Ground</sub> | EEG → CoP | - | Normal<sub>Foam</sub> | CoP → EEG | 2.716 | 150 | 0.207 |
| Narrow<sub>Ground</sub> | EEG → CoP | - | Narrow<sub>Foam</sub> | EEG → CoP | -3.706 | 150 | 0.008 |
| Narrow<sub>Ground</sub> | EEG → CoP | - | Narrow<sub>Foam</sub> | CoP → EEG | 1.803 | 150 | 1 |
| Narrow<sub>Ground</sub> | CoP → EEG | - | Normal<sub>Foam</sub> | CoP → EEG | -0.816 | 150 | 1 |
| Narrow<sub>Ground</sub> | CoP → EEG | - | Narrow<sub>Foam</sub> | CoP → EEG | -1.686 | 150 | 1 |
| Normal<sub>Foam</sub> | EEG → CoP | - | Normal<sub>Ground</sub> | CoP → EEG | 7.622 | 150 | < .001 |
| Normal<sub>Foam</sub> | EEG → CoP | - | Narrow<sub>Ground</sub> | CoP → EEG | 7.508 | 150 | < .001 |
| **Normal<sub>Foam</sub>** | **EEG → CoP** | - | **Normal<sub>Foam</sub>** | **CoP → EEG** | **6.692** | **150** | **< .001** |
| Normal<sub>Foam</sub> | EEG → CoP | - | Narrow<sub>Foam</sub> | EEG → CoP | 0.22 | 150 | 1 |
| Normal<sub>Foam</sub> | EEG → CoP | - | Narrow<sub>Foam</sub> | CoP → EEG | 5.729 | 150 | < .001 |
| Normal<sub>Foam</sub> | CoP → EEG | - | Narrow<sub>Foam</sub> | CoP → EEG | -0.879 | 150 | 1 |
| Narrow<sub>Foam</sub> | EEG → CoP | - | Normal<sub>Ground</sub> | CoP → EEG | 7.31 | 151 | < .001 |
| Narrow<sub>Foam</sub> | EEG → CoP | - | Narrow<sub>Ground</sub> | CoP → EEG | 7.194 | 150 | < .001 |
| Narrow<sub>Foam</sub> | EEG → CoP | - | Normal<sub>Foam</sub> | CoP → EEG | 6.388 | 150 | < .001 |
| **Narrow<sub>Foam</sub>** | **EEG → CoP** | - | **Narrow<sub>Foam</sub>** | **CoP → EEG** | **5.455** | **150** | **< .001** |

**Figure 4:**
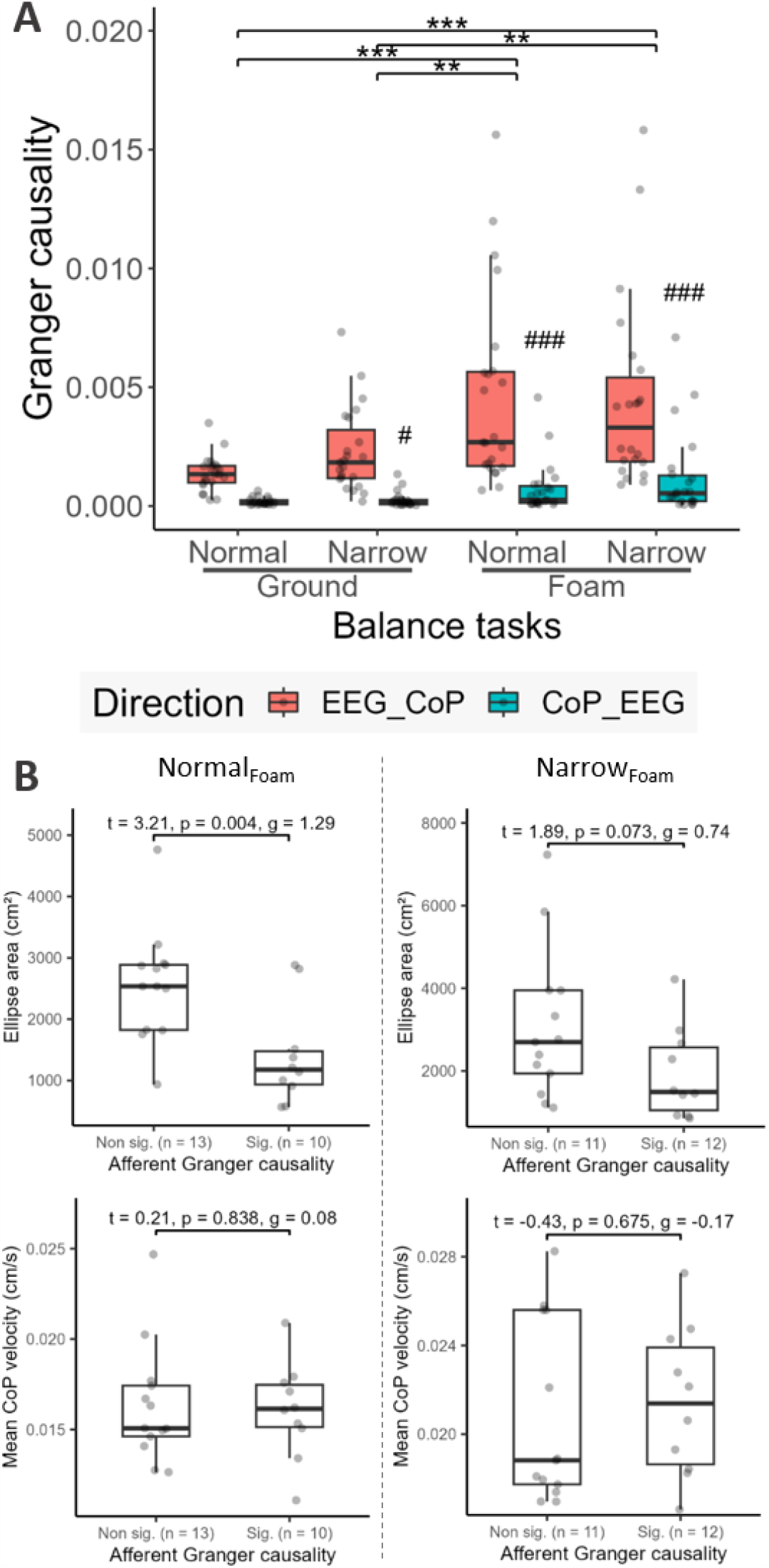
**A:** Time domain GC between the CoP velocity and the Cz electrode in the EEG to CoP velocity (red) and velocity of the CoP velocity to EEG (blue) information flows (upper panel). **B:** 95% confidence ellipse area (top right) and mean CoP velocity (bottom right) differences between participants not showing (GC^-^) and showing (GC^+^) significant GC in the CoP velocity to EEG information flow for the Normal_Foam_ (lower panel, left column) and Narrow_Foam_ (lower panel, right column) conditions. Dots in boxplots represent participants’ individual values. #: p < .05, ##: p < .01, ###: p < .001. *: p < .05, **: p < .01, ***: p < .001.

All participants showed significant GC in the EEG to CoP velocity direction in each experimental condition. Conversely, in the CoP velocity to EEG direction, no participants showed significant GC in Normal_Ground_ and Narrow_Ground_ conditions, and 10 and 12 participants showed significant GC in Normal_Foam_ and Narrow_Foam_ conditions, respectively. Based on these observations, we compared the 95% confidence ellipse area and the mean CoP velocity between participants showing (GC^+^) and those not showing (GC^-^) GC in the CoP velocity to EEG direction for the Normal_Foam_ and Narrow_Foam_ conditions using linear models (Figure 4, lower panels). Model results are presented with r-squared estimates, t values, degrees of freedom, p-values and effect size. The results revealed a significant greater area of the 95% confidence ellipse (1168 cm^2^ comprised in the 95% confidence interval [400 1935]) in the GC^-^ than in the GC^+^ group (r^2^ = 0.29, t_21_ = 3.21, p = 0.004, g = 1.29) in the Normal_Foam_ condition. In the Narrow_Foam_ condition, the area of the 95% confidence ellipse tended to be greater in the GC^-^ than the GC^+^ groups (r^2^ = 0.08, t_21_ = 1.89, p = 0.07, g = 0.74). In both groups, the mean CoP velocity did not significantly differ.

## 4 Discussion

In this study, we aimed to investigate the sensorimotor cortex involvement in postural control during quiet standing tasks with various levels of difficulty in healthy young adults. As the postural task difficulty increased, synchronization between cortical and CoP velocity oscillations in the Delta and Theta frequency bands increased. Moreover, our results showed a correlation between postural performance and this cortico-to-postural coherence in the most challenging postural tasks. Contrary to our initial hypothesis (Bourguignon et al., 2015, 2019), GC analyses unveiled that this cortico-to-postural synchronization mostly results from an efferent information flow. Our study is the first experimental evidence of the control of the CoP velocity by the central nervous system through coherence and effective connectivity analyses. Interestingly, participants with reduced afferent contributions between cortical and CoP velocity oscillations had reduced postural performance, suggesting that postural control relies on both feedforward and feedback mechanisms.

### 4.1 Validation of the balance disturbance protocol

From a behavioural perspective, the alteration of postural performance during quiet standing has been commonly investigated through experimental paradigms employing various stance width (Bonnet & Despretz, 2012), surface compliance (Patel et al., 2008), or their combination (Liang et al., 2022). The analysis of the 95% confidence ellipse area and the mean CoP velocity revealed a decreased postural performance during tasks of increasing difficulty. Hence, our results showed that our experimental manipulation successfully altered postural control by combining various stance and compliant ground strategies (Doyle et al., 2005; Quijoux et al., 2021). These results show the relevance of our experimental paradigm to study cortical involvement during postural task of increasing difficulty. As to the power spectrum of the CoP velocity, Demura et al. (2008) previously reported that in healthy young adults during quiet standing, 75% and 99% of the total power spectrum of CoP velocity was comprised in the 0-2 Hz frequency band and below 6 Hz, respectively. In line with these previous findings, the CoP velocity power spectrum in our study showed that 99% of the total power spectrum was comprised within the 0-8 Hz frequency band, with 52%-57% in the Delta band and 11%-15% in the Theta band depending on the experimental conditions. In addition to justifying the frequency bands selected for the connectivity analyses, the spectral power results confirm the decreased stability associated with the increase of task difficulty.

### 4.2 Power spectra patterns during quiet standing

Previous studies investigating cortical activity modifications during balance perturbation paradigms consistently showed changes in Beta, Alpha, Theta, and Delta frequency bands power spectrum. Alpha and Beta bands desynchronization was observed in the centroparietal area (Edwards et al., 2018; Hülsdünker et al., 2016), primarily linked to sensory information processing during quiet standing. While not significantly different from baseline, our results showed a similar pattern in the Beta band TRSP over the central area and in the Alpha band in the centroparietal area during the most challenging experimental condition. The increased Theta band spectral power in the central area has previously been observed using external perturbations paradigms (Sipp et al., 2013; Varghese et al., 2014) or during quiet standing in highly challenging conditions, such as sensory input modification with eyes closed (Edwards et al., 2018) or a smaller base of support with unipedal stance (Hülsdünker et al., 2015, 2016). Again, our results showed a similar, although not significant, pattern in Theta band TRSP during the most challenging experimental condition. Additionally, one previous study reported increased spectral power in the Delta band in the central area, a finding that was reproduced in our study’s most challenging experimental condition (Ozdemir et al., 2018).

The absence of statistically significant differences in TRSP results across our experimental conditions may be attributed to two key factors. First, previous studies did not use an experimental reference condition (Edwards et al., 2018; Hülsdünker et al., 2015, 2016), which can contribute to discrepancies in observed outcomes. Second, our experimental paradigm entailed simpler tasks compared to those previously used in the existing literature. This discrepancy arises from the absence of sensory input manipulation with closed eyes (Edwards et al., 2018; Ozdemir et al., 2018), the use of narrower base of support (Hülsdünker et al., 2015, 2016), or the introduction of mechanical perturbations (Sipp et al., 2013; Slobounov et al., 2009). Nevertheless, the consistent patterns of postural performance alteration with increasing task difficulty, along with cortical activity profiles that align with previous findings, support the assertion that our experimental paradigm effectively challenges postural control and elicits changes in cortical activity. To further investigate cortical contribution to quiet standing control, a more comprehensive approach is needed. Indeed, maintaining an upright posture requires continuous corrective actions for maintaining the CoP within the base of support. Therefore, extracting events from the CoP velocity and investigating the corresponding brain dynamics could be a solution to provide a better understanding of cortical involvement during postural control (Stins & Roerdink, 2018).

### 4.3 Corticopostural coherence

The first investigations of the coupling between brain oscillations and body kinematics showed a synchronization between cortical oscillations from the sensorimotor motor cortex area and movement velocity in low frequency band (2-5 Hz) (Jerbi et al., 2007; Kelso et al., 1998). Subsequent studies extended these results to movement acceleration during repetitive finger movements (Bourguignon et al., 2011, 2012). However, to the best of our knowledge, the present study is the first to evidence a synchronization between cortical oscillations and postural kinematics using coherence and effective connectivity analyses. Our findings revealed a significant synchronization in the Delta and Theta frequency bands between the CoP velocity and cortical oscillations recorded at the Cz electrode site and neighbouring electrodes. This central coherence peak is in line with previous results (Bourguignon et al., 2012; Jerbi et al., 2007) and is consistent with the synchronization between the primary motor or somatosensory areas and body kinematics previously reported (Bourguignon et al., 2019). Additionally and according to the literature on CKC, the maximum coherence values were observed within the Delta band (Bourguignon et al., 2019; Jerbi et al., 2007; Kelso et al., 1998), emphasizing the importance of this frequency band in sensorimotor synchronization during postural control. Moreover, the increased coherence in the Delta band between cortical oscillations and CoP velocity could reflect a greater communication between the sensorimotor cortex and postural outcomes when individuals confront balance challenges during quiet standing.

Previous works showed the significant role of body sway velocity in postural control (Masani et al., 2003, 2014). These studies suggested that postural control relies on velocity information to activate muscles in anticipation of body sway (Masani et al., 2003). Moreover, models of postural control based on CoP velocity outperformed models relying on CoP position for predicting body CoM displacements (Delignières et al., 2011). Our results agree with the importance of CoP velocity in postural control, offering physiological evidence that establishes a communication between brain oscillations and kinematic outcomes. Hence, given both the synchronization between brain and CoP velocity oscillations, and by extension body sway velocity (Richmond et al., 2021), during postural tasks and the existing models of postural control found on velocity information, we propose to name this coupling corticopostural coherence. This term extends the concept of corticokinematic coherence previously introduced (Bourguignon et al., 2019). Taken together, our results contribute to further evidence the sensorimotor cortex involvement when balance is challenged during quiet standing.

To go beyond the results provided by coherence analysis that lack information flow direction, GC was used to deepen our understanding of corticopostural coherence mechanism(Bastos & Schoffelen, 2016). Earlier studies revealed that CKC mainly results from afferent mechanisms (Bourguignon et al., 2015, 2017; Piitulainen et al., 2013), which were interpreted as the integration of proprioceptive information of peripheral feedback (Bourguignon et al., 2019). However, our results showed a higher time domain GC in the efferent compared to the afferent direction. Given that postural control relies on multisensory information integration (Maurer et al., 2006; Mergner et al., 2002) and, notably, proprioceptive information (Jeka et al., 1998), our results suggest that corticopostural coherence reflects a distinct mechanism from CKC. Interestingly, previous studies showed a prevalence of efferent information between EEG and electromyography signals during quiet standing (Liang et al., 2022) and a positive time lag between the activity of ankle and knee muscles and CoP displacements (Gatev et al., 1999). This leads to the interpretation that ankle muscles are controlled by the motor cortex in an anticipatory planning process during quiet standing. Consequently, corticopostural coherence could reflect a feedforward mechanism during postural tasks (Masani et al., 2003; Stins & Roerdink, 2018) involving the sensorimotor cortex through the synchronization of cortical oscillations and CoP velocity to maintain balance. As discussed in the subsequent section, this corticopostural coherence mechanism seems crucial for maintaining balance in challenging tasks.

### 4.4 Importance of the corticopostural coherence mechanism in postural control

As previously discussed, cortical activity in Theta and Delta frequency bands scales with task difficulty during quiet standing (Hülsdunker, 2015, 2016; Edwards, 2018; Ozdemir, 2018), emphasizing the increased sensorimotor cortex involvement in postural control. Our results showing increased Delta corticopostural coherence in difficult tasks provides further insights into the sensorimotor cortex’s role in balance maintenance. Moreover, the negative correlation between Delta corticopostural coherence and the area of the 95% confidence ellipse in the most challenging condition suggests that participants with the highest coherence values were more stable. Therefore, corticopostural coherence may constitute a fundamental mechanism involved when postural control is challenged.

From the analysis of the direction of information flow between the sensorimotor cortex and postural outcomes, our results showed a significant effect of postural task difficulty on GC and an interaction between task difficulty and information flow direction, leading to a greater difference between afferent and efferent directions in the most challenging tasks. Therefore, the increased corticopostural coherence seems mainly driven by an increased efferent information flow conveyed by the sensorimotor cortex. One might hypothesize that the central nervous system exerts an increased control of the CoP velocity through increased synchronization between brain oscillations and CoP velocity during difficult postural tasks (Fries, 2015; Richmond et al., 2021). Furthermore, our results also showed significant afferent GC in 10 and 12 out of the 23 participants in the Normal_Foam_ and Narrow_Foam_ conditions, respectively, and that for those participants postural performance was better. In accordance with the interpretation of CKC, which is mainly driven by sensory information integration (Bourguignon et al., 2019), our results emphasize the critical role of CoP velocity information integration in maintaining balance during challenging tasks. Thus, our combined results suggest the involvement of both feedforward and feedback mechanisms in postural control when facing challenging conditions (Fitzpatrick et al., 1996; Stins & Roerdink, 2018). It’s important to note that GC analyses do not establish definitive causality between two oscillating signals but rather provide insights into the sequence of one signal compared to another (Bastos & Schoffelen, 2016). Therefore, the interpretation of our results should be limited to understanding the direction of information flow within the brain networks under investigation. To confirm these results, further research should aim to confirm the GC findings by altering CoP velocity information.

### 4.5 Limitations

The spatial resolution of cortical activity with a 64-electrode EEG recording proves inadequate for distinguishing between the primary motor area and the primary somatosensory area (Bourguignon et al., 2019; Edwards et al., 2018). Consequently, further exploration of the origins of the observed brain oscillations necessitates increasing spatial resolution using denser EEG caps and implementing more advanced processing techniques, such as source reconstruction. Regarding the postural tasks, it is essential to note that the paradigm used in this study involved exclusively quiet standing. Therefore, the applicability of these results to real-life situations may be limited since quiet standing does not involve external perturbations and fails to encompass all the challenges encountered in daily activities (Visser et al., 2008). Nonetheless, previous studies have established a correlation between the risk of falls in older adults and performance in quiet standing tasks (Quijoux et al., 2020), indicating that mechanism of the postural control can transfer effectively between quiet standing paradigms and daily-living situations.

### 4.6 Conclusion

This study is the first to reveal significant coherence between oscillations from the central sensorimotor cortex and CoP velocity during quiet standing, thus extending the CKC to postural control, herein named corticopostural coherence. The analysis of the information flow direction revealed a dominant efferent information transfer, confirming the central role of CoP velocity in postural control. Additionally, these results are further supported by positive correlations between postural performance and corticopostural coherence and augmented postural performance in participants showing significant afferent GC in challenging postural task, reflecting the importance of this mechanism to maintain quiet stance.

