## Supplementary material for "Corticopostural functional and effective connectivity reveal cortical control of postural sway velocity during quiet standing"

Supplementary Materials

3.1. Postural outcomes: ellipse area model

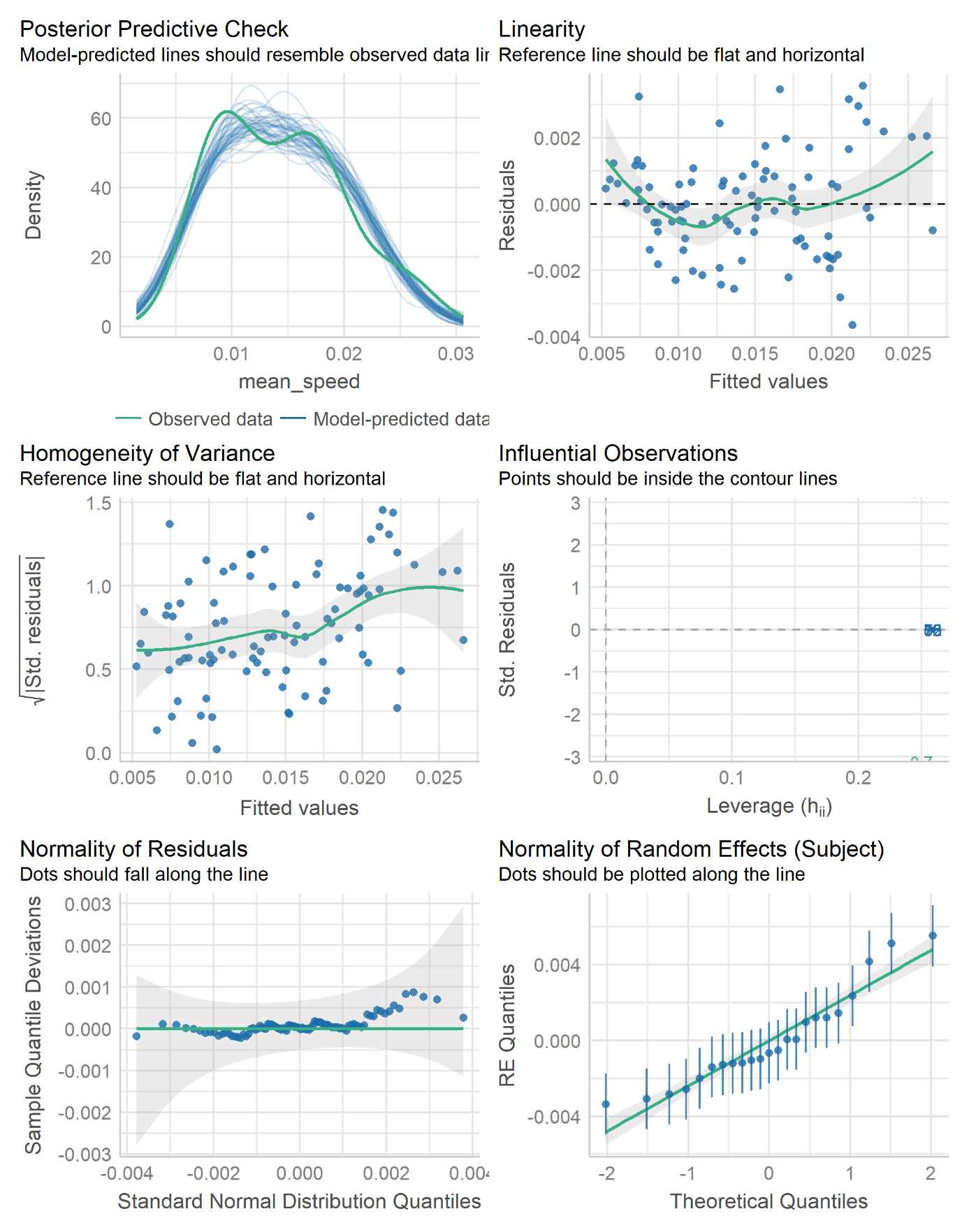

3.1. Postural outcomes: CoP mean speed model

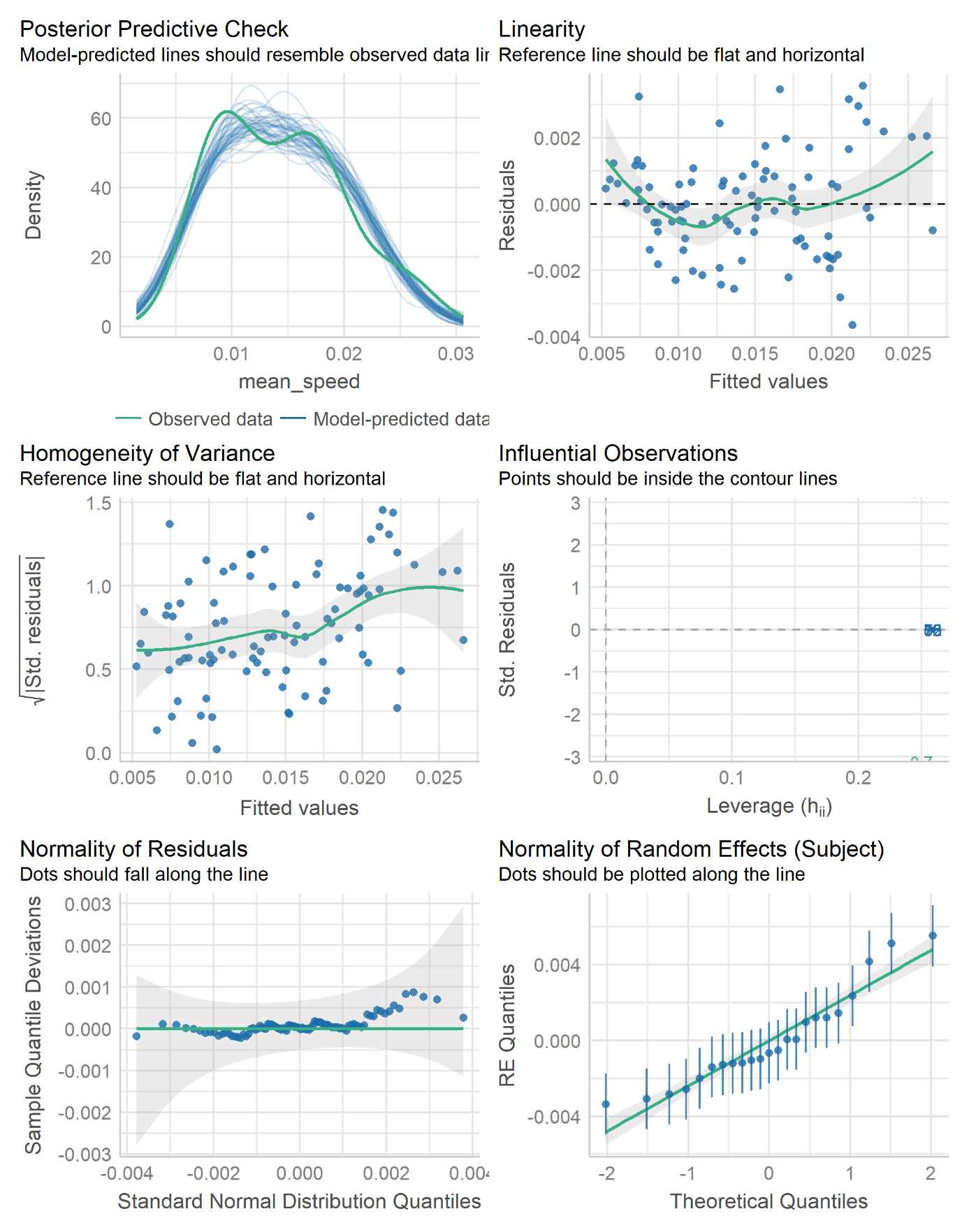

3.3. Coherence between cortical and CoP velocity oscillations: Theta band coherence’s model

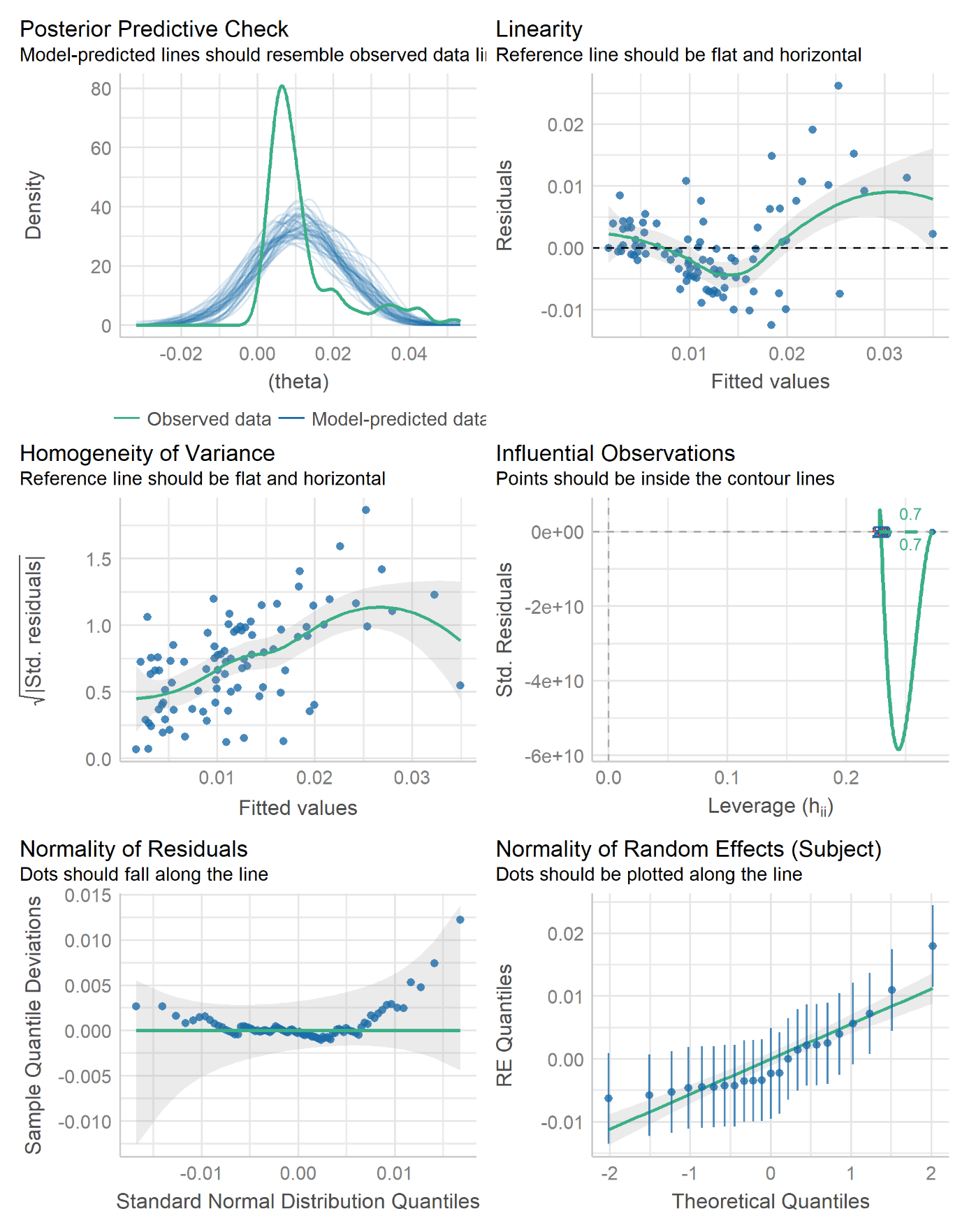

3.3. Coherence between cortical and CoP velocity oscillations: Delta band coherence’s model

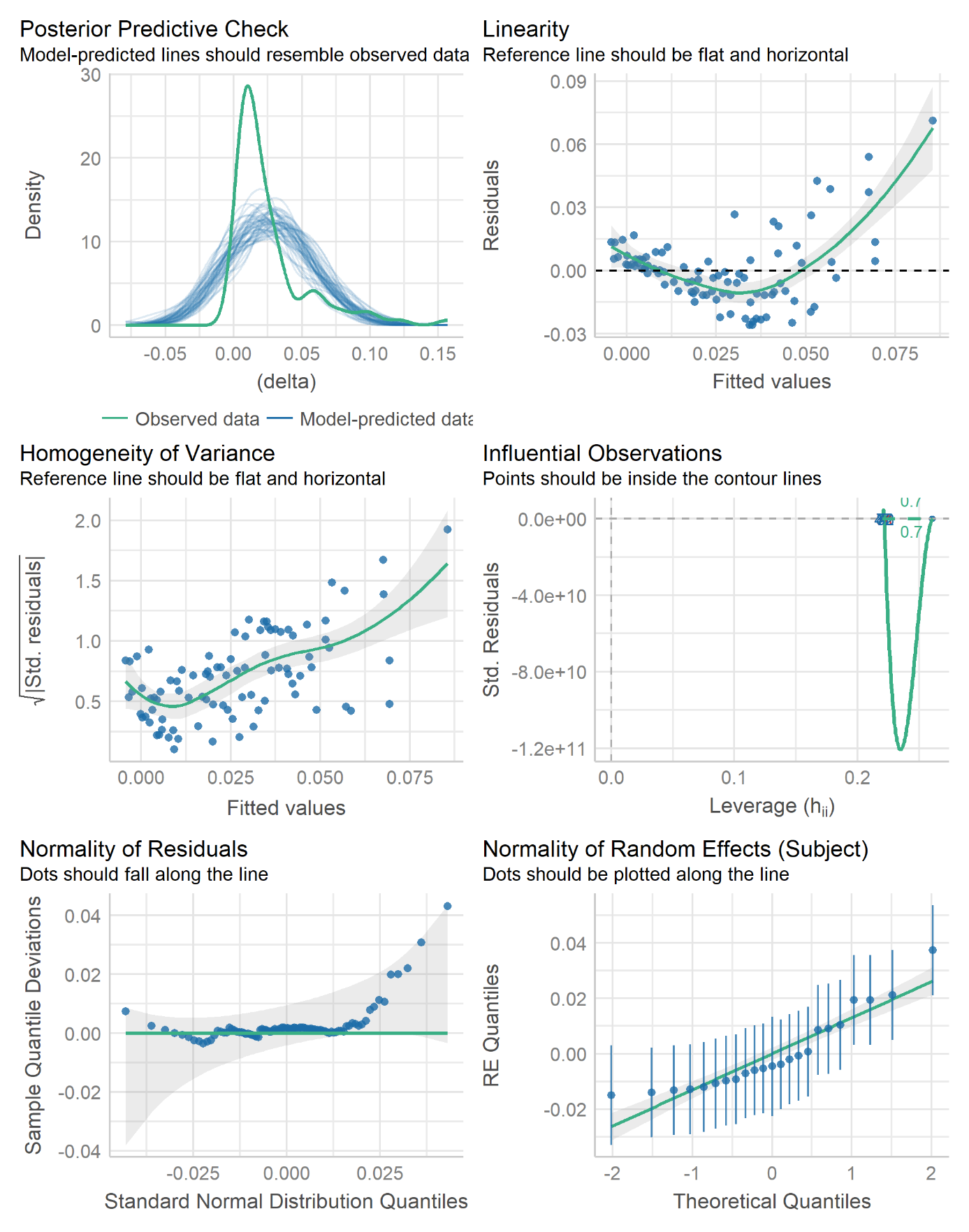

3.4. Granger causality between cortical and CoP velocity oscillations: GC’s model

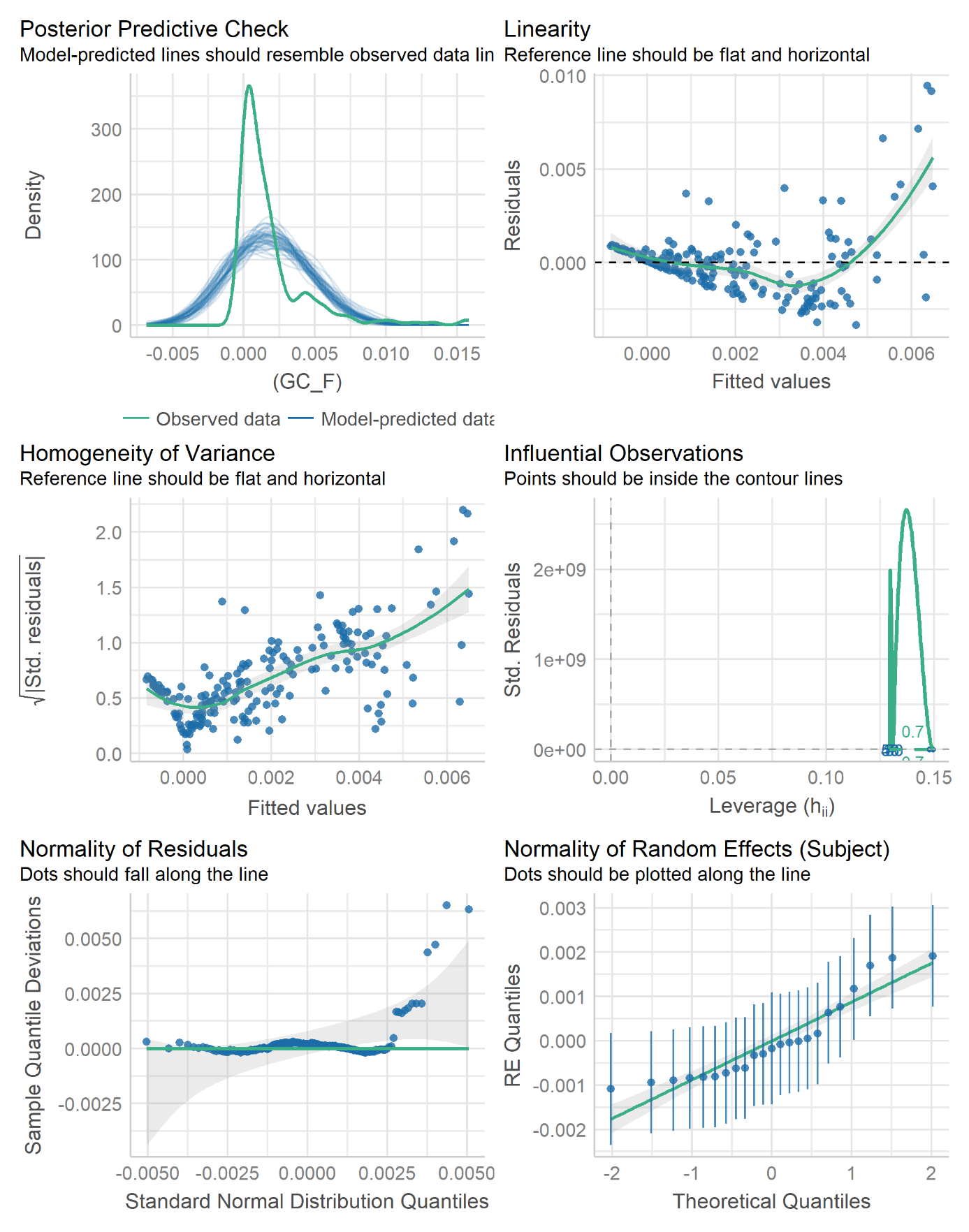
